# Expanding the Molecular Alphabet of DNA-Based Data Storage Systems with Neural Network Nanopore Readout Processing

**DOI:** 10.1101/2021.09.27.462049

**Authors:** S. Kasra Tabatabaei, Bach Pham, Chao Pan, Jingqian Liu, Shubham Chandak, Spencer A. Shorkey, Alvaro G. Hernandez, Aleksei Aksimentiev, Min Chen, Charles M. Schroeder, Olgica Milenkovic

## Abstract

DNA is a promising next-generation data storage medium, but the recording latency and synthesis cost of oligos using the four natural nucleotides remain high. Here, we describe an improved DNA-based storage system that uses an extended 11-letter molecular alphabet combining natural and chemically modified nucleotides. Our extended-alphabet molecular storage paradigm offers a nearly two-fold increase in storage density and potentially the same order of reduction in the recording time. Experimental results involving a library of 77 custom-designed hybrid sequences reveal that one can readily detect and discriminate different combinations and orders of monomers via MspA nanopores. Furthermore, a neural network architecture designed to classify raw current signals generated by Oxford Nanopore Technologies sequencing ensures an average accuracy exceeding 60%, which is 39 times higher than that of random guessing. Molecular dynamics simulations reveal that the majority of modified nucleotides do not induce dramatic disruption of the DNA double helix, making the extended alphabet system potentially compatible with PCR-based random access data retrieval. The methodologies proposed provide a forward path for new implementations of molecular recorders.

---

DNA is emerging as a data storage medium that offers ultrahigh storage density and a level of robustness not matched by conventional magnetic and optical recorders. Information stored in DNA can be copied in a massively parallel manner and selectively retrieved via polymerase chain reaction (PCR) (1–8). However, existing DNA storage systems suffer from high latency caused by the inherently sequential writing process. Despite recent progress, a typical cycle time of solid-phase DNA synthesis is on the order of minutes, which limits practical applications of this molecular storage platform (9). Using current technologies, writing 100 bits of information (or, roughly two words in this article) requires nearly two hours and costs more than US$1, assuming that each nucleotide stores its theoretical maximum of two bits. To overcome these and other challenges, new synthesis methods and/or new information encoding approaches are required to accelerate the speed of writing large-volume data sets (10).

Expanding the alphabet of a DNA storage media by including chemically modified DNA nucleotides can both increase the storage density and the writing speed as more than two bits are recorded during each synthesis cycle. However, designing chemically modified DNA nucleotides as new letters for the DNA storage alphabet must be tightly coupled to the process of reading the encoded information, i.e., DNA sequencing, because current DNA sequencing methods, including nanopore sequencing, have been developed and optimized to read biological nucleotides. Prior work reported an expanded nucleic acid alphabet of eight synthetic DNA and RNA nucleotides that can be replicated and transcribed using biological enzymes (11). That alphabet, however, was not designed for molecular storage applications and has not been read accurately using a nucleic acid sequencing method. Aerolysin nanopores were used to detect synthetic polymers flanked by adenosines, where each monomer of the polymer carries one bit of information (12). A proof-of-principle study has also shown that a base pair containing a chemically modified nucleotide could be replicated and read using a biological nanopores (13).

Here, we report on an expanded molecular alphabet for DNA-based data storage comprising four natural and seven chemically modified nucleotides (**Table 1**, **Figures 1, S1-S3**) that are readily detected and distinguished using nanopore sequencers. Our results show that MspA nanopores can accurately discriminate 77 diverse combinations and orderings of monomers within homo- and heterotetrameric sequences (as listed in **Tables S2-S4**).

**Figure 1.**
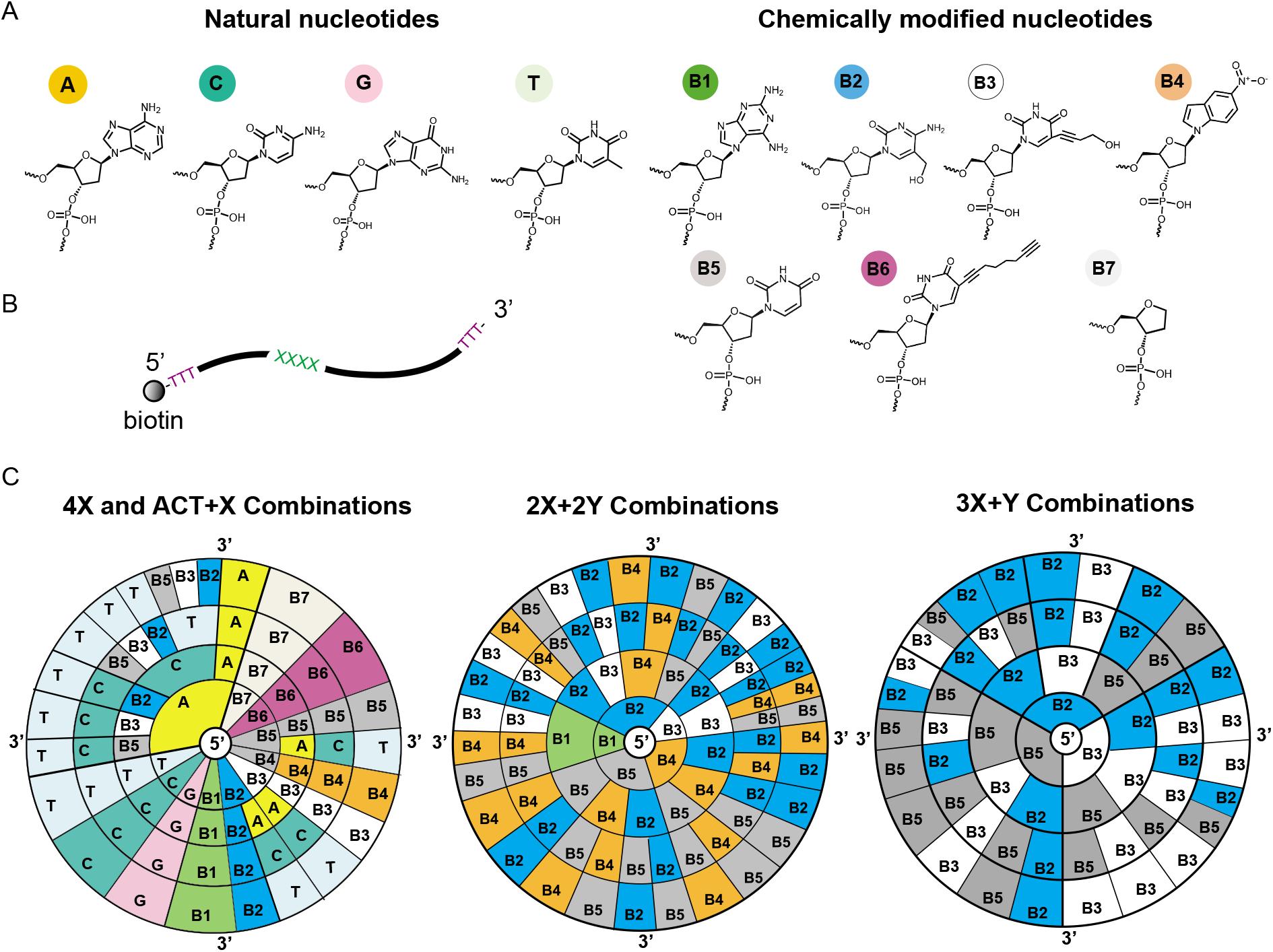
DNA data storage using natural and chemically modified nucleotides. **(A)** Chemical structures of natural DNA nucleotides (A, C, G, T) and the selected chemically modified nucleotides employed in our study (B1-B7). **(B)** Schematic of the ssDNA oligo organization used in MspA nanopore experiments. The length of the oligos is 40 nucleotides (nts), with biotin attached at the 5’ terminus. Homo- or heterotetrameric sequences are located at positions 13-16, flanked by two polyT regions of length 12 nt and 24 nt on the 5’ and 3’ ends, respectively. **(C)** The sequence space for DNA homotetramers or heterotetramers used in the MspA nanopore experiments. The notation aX+bY, where a and b take values in {2,3,4} so that a + b = 4, indicates that ‘a’ symbols of the same kind are combined with ‘b’ symbols of another kind, and arranged in an arbitrary linear order. In total, 77 distinct tetrameric sequences were synthesized and tested experimentally. (Left) Circular diagram showing all 11 homotetramers and 12 tetrameric sequences of the form ACT+X, where X is a chemically modified nucleotide from the set {B2, B3, B5}. (Middle) Circular diagram showing all 30 tested combinations of tetrameric sequences with total composition 2X+2Y using chemically modified monomers from the set {B1, B2, B3, B4, B5}, including sequence patterns XXYY, XYYX, and XYXY. (Right) Circular diagram showing the remaining 24 combinations of tetrameric sequences with total composition 3X+Y using the set {B2, B3, B5}. Five chemically modified nucleotides form stable base pairs with natural nucleotides via hydrogen bonds (B2—G, B3—A, B5—A, B6—A, B6— C), based on the results from molecular dynamic (MD) simulations.

**Table 1.**
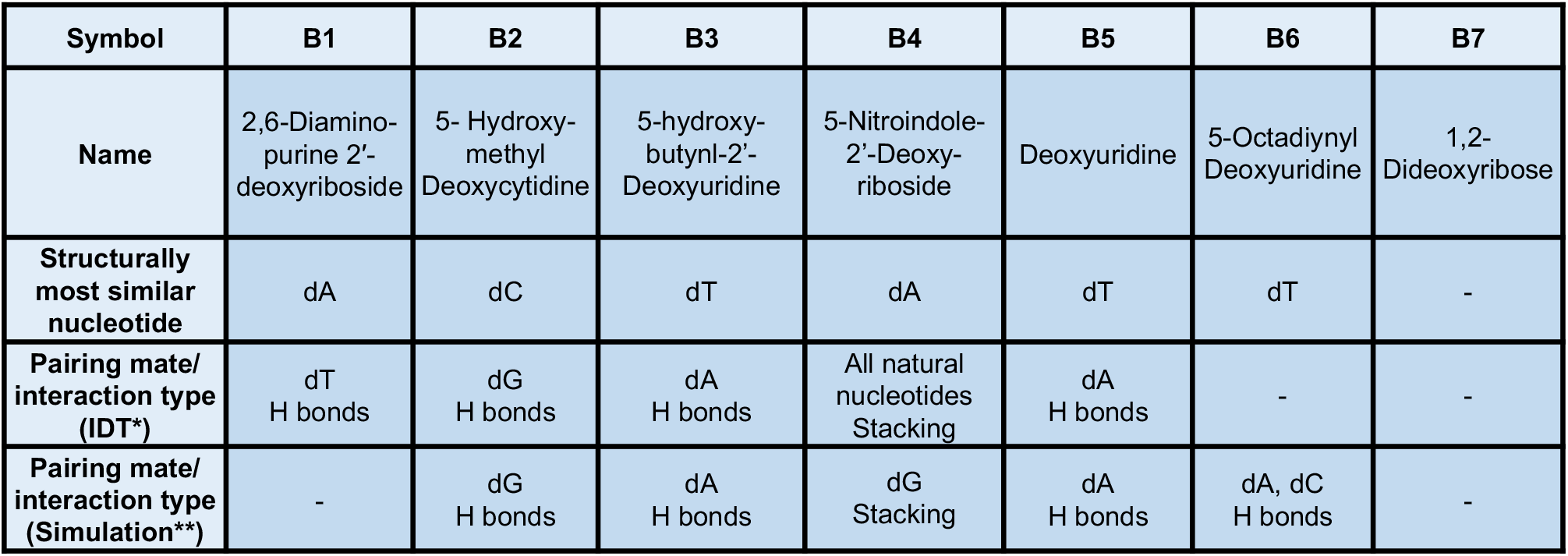
Chemically modified nucleotides used in the proposed DNA data storage system, along with their chemical properties. The symbols and the names of the chemically modified nucleotides are shown in the first and second row, while the molecular structures are depicted in Figure 1. Structurally similar natural nucleotides are shown in the third row. In general, distinct chemical functional groups and molecular charges play an important role in discriminating monomers using MspA and ONT sequencers. The last two rows show pairing properties of the modified bases: * denotes data from Integrated DNA Technologies (14) while ** denotes results from molecular dynamics simulations reported in the Supplementary Information (**Figure S1-2**, **Table S1**). Short dashes indicate that pairing is inherently impossible (e.g., B7) or that no stable interactions were identified.

| Symbol | B1 | B2 | B3 | B4 | B5 | B6 | B7 |
| --- | --- | --- | --- | --- | --- | --- | --- |
| Name | 2,6-Diamino-purine 2'-deoxyriboside | 5- Hydroxy-methyl Deoxycytidine | 5-hydroxy-butynl-2'-Deoxyuridine | 5-Nitroindole-2'-Deoxy-riboside | Deoxyuridine | 5-Octadiynyl Deoxyuridine | 1,2-Dideoxyribose |
| Structurally most similar nucleotide | dA | dC | dT | dA | dT | dT | - |
| Pairing mate/ interaction type (IDT*) | dT<br>H bonds | dG<br>H bonds | dA<br>H bonds | All natural nucleotides Stacking | dA<br>H bonds | - | - |
| Pairing mate/ interaction type (Simulation**) | - | dG<br>H bonds | dA<br>H bonds | dG Stacking | dA<br>H bonds | dA, dC<br>H bonds | - |

We further demonstrate that highly accurate classification (exceeding 60% on average) of combinatorial patterns of natural and chemically modified nucleotides is possible using deep learning architectures that operate on raw current signals generated by GridION of Oxford Nanopore Technologies (ONT).

Stable bonding of chemically modified nucleotides within a DNA double helix is important for DNA-based storage because it enables durable preservation of recorded information, as well as random access to the stored data by means of PCR reactions (4). To better understand the interactions between chemically modified and natural nucleotides, we also investigated the stability of modified DNA duplexes by carrying out all-atom molecular dynamics (MD) simulations of the Dickerson dodecamers (15) containing a pair of chemically modified nucleotides (**Figure 2A**). Out of many possible variants, we chose to investigate the stability of B1—T, B2—G, B3—A, and B5—A base pairs, as suggested by Integrated DNA Technologies (IDT), as well as the pairing of B4 and B6 with all four types of natural nucleotides. Each modified dodecamer was solvated in electrolyte solution and simulated for approximately 350 ns. Five modified-natural base pairs, (B2—G, B3—A, B5— A, B6—A, and B6—C) were found to form stable hydrogen bond patterns within the duplex forming either two or three hydrogen bonds per base pairs (**Figure 2B-E**). The average number of hydrogen bonds was found to be 1.37 for B2—G, 1.01 for B3—A, 1.00 for B5— A, 1.00 for B6—A and 0.70 for B6—C, which are results compatible with the numbers computed for the canonical base pairs (0.83 for A—T and 1.23 for C—G) using the same hydrogen bond criteria. In all other modified-natural combinations, we observed local disruptions of the base pairing structure (**Figures S1-2**). In B1—T, B4—A and B4—T pairs, the bases were observed to protrude out from the duplex without disrupting the hydrogen bonding of the surrounding base pairs. The B6—G pair formed a base stacking pattern, forcing the breakage of hydrogen bonds in the adjacent base pairs. Local unraveling of the duplex structure was observed in the systems containing B4—G, B4—C and B6—T base pairs. Based on these results, we conclude that most of our chemically modified nucleotides introduce minor perturbations to the structure of the duplex except for B4, which does not fit well within the geometry of the classical DNA duplex but is not sufficient to produce a complete unraveling of the DNA duplex. However, we observed that an isolated B4-G base pair is able to maintain stable stacking interaction when simulated under conditions that mimic the presence of a longer DNA strand (**Figure S2**).

**Figure 2.**
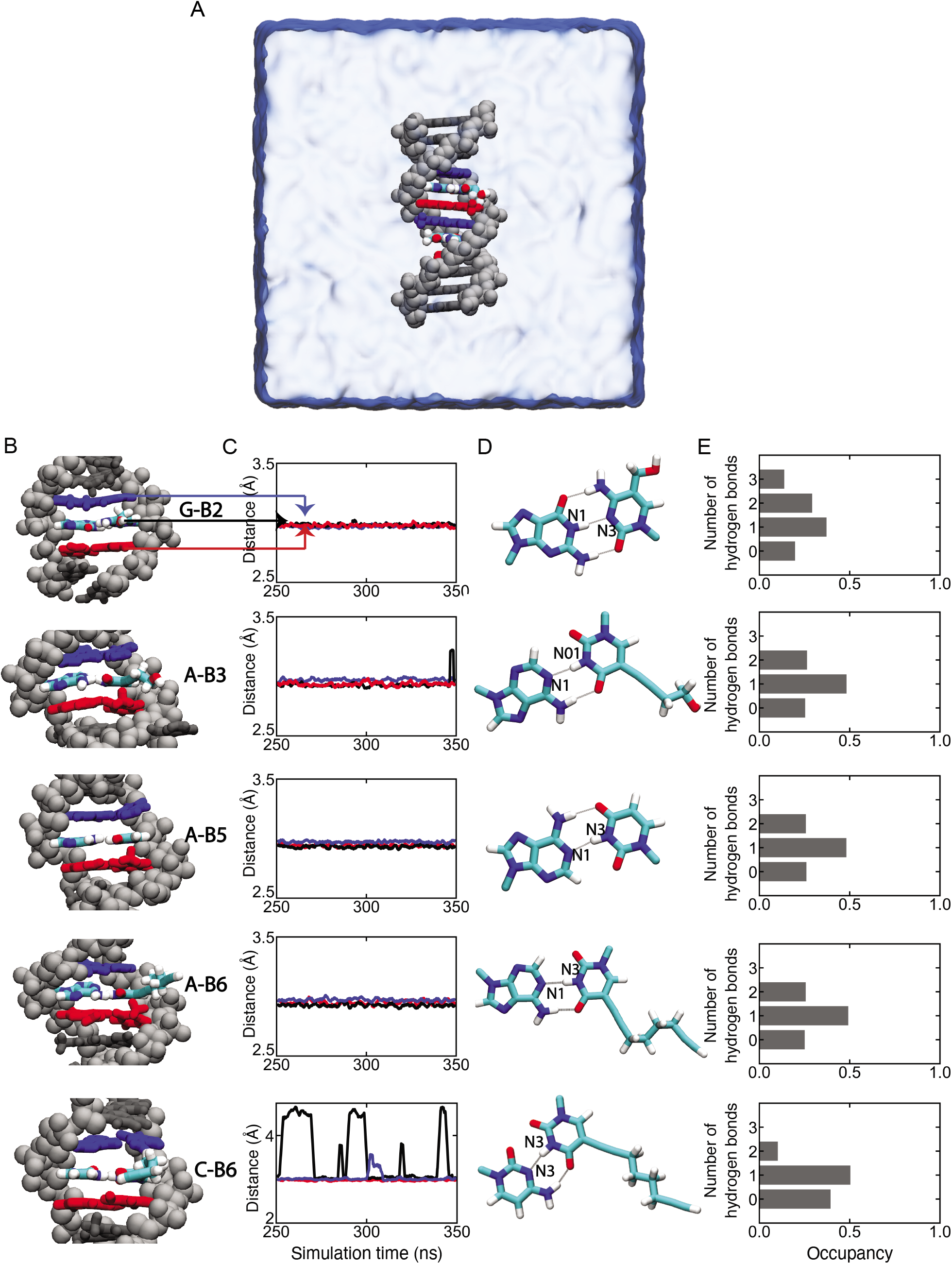
Stability of DNA duplexes containing chemically modified nucleotides. **(A)** Initial state of a simulation system where a DNA dodecamer containing chemically modified nucleotides is immersed in electrolyte solution. The backbone of the dodecamer is shown using silver spheres whereas the bases are drawn as molecular bonds. Chemically modified bases and the natural bases that pair with them are colored according to the atom type (cyan for carbon, blue for nitrogen and red for oxygen). Base pairs immediately adjacent to the modified base pair are colored in red or blue. **(B)** Microscopic configurations of modified base pairs (from top to bottom: B2—G, A—B3, A—B5, A— B6 and C—B6) shown using the same color scheme as in panel B. **(C)** Donor (N1)—acceptor (N3) distance (black) in the modified base pair (black) and in the adjacent base pairs (red and blue) during the last 100 ns of the 350 ns MD simulation. The arrows indicate the correspondence between the base pairs and the curves. The curves show a running average of the 10 ps-sampled data with a 2 ns averaging window. **(D)** Microscopic configuration of modified base pairs. The black lines represent hydrogen bonds. The donor and the acceptor are labeled asides the atoms. **(E)** Probability of observing the specified number of hydrogen bonds within a modified base pair. The H-bonding probabilities were computed using the final 100 ns of a 350 ns all-atom MD simulation of a DNA dodecamer.

To determine whether natural and chemically modified DNA nucleotides can be distinguished by measuring ionic current through biological nanopore MspA, we designed a series of single-stranded DNA (ssDNA) molecules with the general sequence 5’-biotin- (dT)_12_-XXXX-(dT)_24_-3’, where X = {A, T, C, G, B1-B7} (**Figure 3, Figures S3-S4, Tables S2-S4**). We hypothesized that specific chemical modifications to nucleobases such as amines, alkynes, or indole moieties can alter polymer-amino acid interactions in biological nanopores, thereby generating distinct signals in nanopore readouts. In the process, we also took into consideration the stability of base pairing and stacking interactions between natural and chemically modified nucleotides based on the described MD simulations and experiments (**Tables 1 and S1, Figures 2, S1 and S2**).

**Figure 3.**
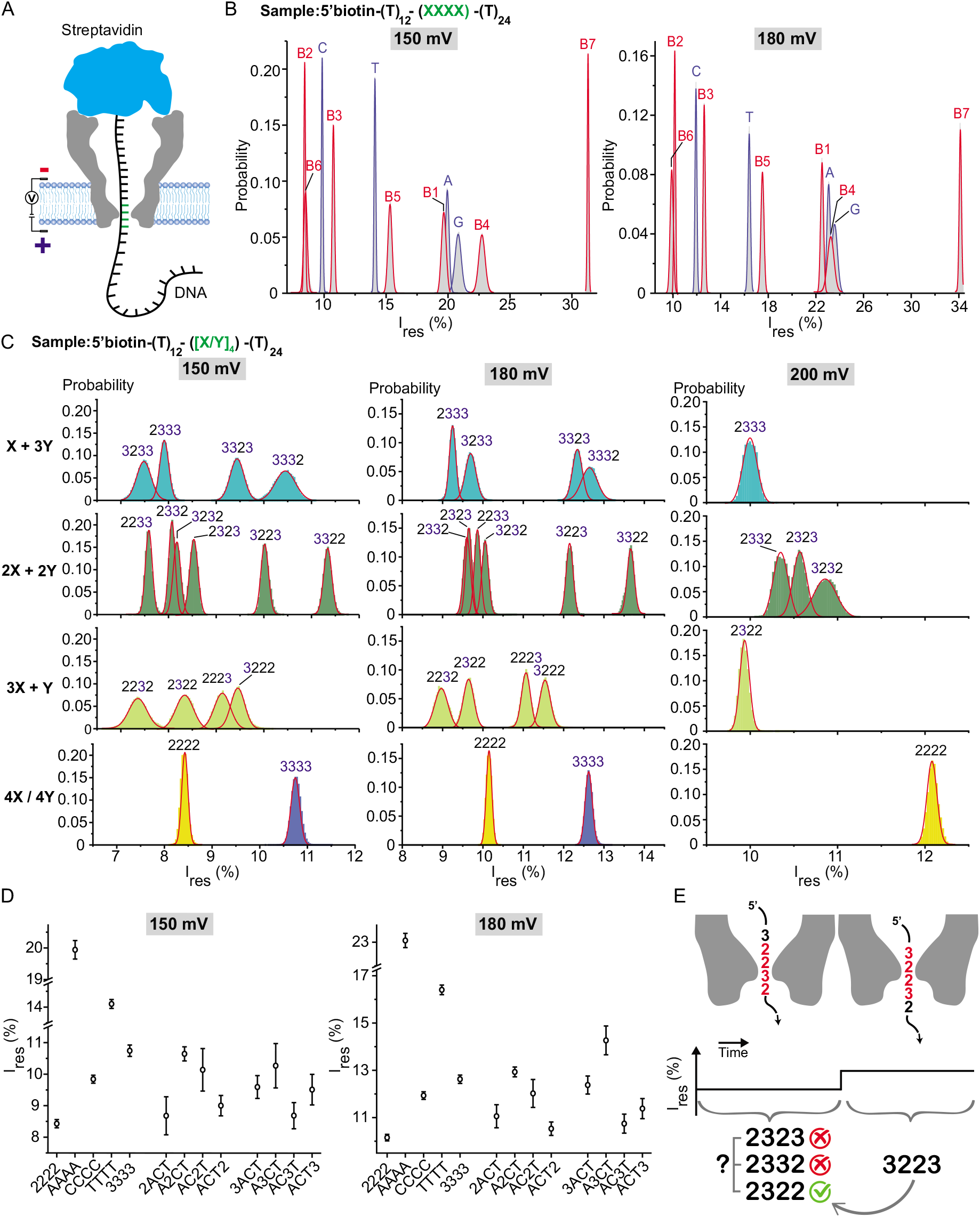
Identification of chemically modified DNA using MspA nanopores. **(A)** Schematic diagram of ssDNA immobilized in a MspA nanopore, where ssDNA containing a biotin-streptavidin interaction at the 5’ terminus prevents translocation through the pore. Residual ion current generated by four nucleotides at positions 13-16 from the 5’ terminus is recorded for ssDNA immobilized in the pore. **(B)** Histograms of average residual ionic currents I_res_ shown in gray for different homopolymers (A, T, C, G, and B1-B7). The fitted Gaussian curves are depicted in red for natural nucleotides (A, T, C, G), and in blue for chemically modified nucleotides (B1-B7). **(C)** Histograms of the average residual ionic currents and the fitted Gaussian curves at various applied voltages for tetramers involving different combinations and orderings of B2 and B3. **(D)** Peak values (points) and confidence intervals (bars) of the fitted Gaussians with mean residual ionic currents corresponding to tetramers obtained by inserting one of the monomers B2 and B3 into the sequence ACT, at applied biases of 150 mV and 180 mV. **(E)** Schematic of the shift reconciliation method for resolving ambiguities in the readouts of different tetramers.

Following molecular design and synthesis of ssDNA oligos, we performed MspA nanopore experiments where ssDNA oligos containing streptavidin at the 5’ terminus were electrophoretically attracted inside MspA nanopores. The bulky streptavidin protein prevents the oligos from fully translocating through the pore without appreciably affecting the measured ionic currents (16). Consequently, ssDNA molecules are effectively immobilized within MspA nanopores, exposing the four nucleotides at positions 13-16 from the tethering point to the constriction of the MspA pore (**Figure 3A**) (17). In this assay, streptavidin holds ssDNA in the MspA constriction similarly to a helicase enzyme that steps through doublestranded (dsDNA) in an ONT sequencer, thereby enabling long duration current readings for each sequence tetramer (**Figure S3**).

We next used MspA nanopores to determine residual currents for homotetrameric sequences of all natural and chemically modified monomers (**Figure 3B**). Our results show that MspA accurately discriminates all four natural (A, G, C, T) and nearly all chemically modified nucleotides (B1-B7) at an applied bias of 150 mV. The abasic nucleotide B7 shows the largest residual current, which likely arises due to its small molecular size and reduced ability to interact with the reading head of MspA. The residual current levels are sensitive to the chemical identity of the nucleotides but do not directly correlate with their molecular size (**Figure 3B**). For example, current signals from B6 and B2 overlap at 150 mV, but B6 is well separated from B3 despite being structurally similar. We further studied the effect of the applied bias on the resolution of nucleotide bases. At 150 mV, four chemically modified nucleotides (B2, B3, B4, B5) showed well-resolved signals from each other and the natural nucleotides, but the current levels from B6 exhibited some overlap with B2. Upon increasing the applied bias to 180 mV, B6 was readily resolved from B2. In addition, at 180 mV, resolution in the I_res_ region exceeding 20% decreased, as may be seen from the residual currents of B4, A, and G which have Gaussian readout distributions which overlap in area by more than 90% (**Figure 3B**).

We further used MspA to detect and identify heterotetrameric sequences with compositions 2X+2Y, where X, Y = {B2, B3, B4, B5} (**Figure 3C**, **Figures S2**-**S3**, **Tables S2**-**S4**). Our results show that MspA can distinguish all heterotetrameric sequences with the same nucleotide composition when measurements at all three applied biases (150 mV, 180 mV, 200 mV) are performed. Due to the large sequence space explored, here we focus our discussion on representative tetrameric combinations of B2 and B3 (**Figure 3C**). In most cases, the residual currents of heterotetramers fall between those of two corresponding homotetramers. For example, the tetramer 3223 has an Ires of 12.3%, whereas those of B2 and B3 are 10.2% and 12.6%, respectively (at 180 mV). However, some combinations of B2 and B3, including 2232, 2322, 2333, 3233, 2323, 2332, and 2233, showed significant decreases in residual currents compared to homotetramers B2 and B3 (**Figure 3C**), whereas the residual current of tetramer 3322 is larger than homotetramers of B2 and B2 at either 150 mV or 180 mV. Importantly, all tetrameric sequences were resolved by adjusting the applied bias (18). At a higher applied bias of 200 mV, tetramers that were unresolved at lower bias were readily resolved, including 2322, 2332, and 2322 (**Figure 3C**). Overall, these results are consistent with the observation that the residual current levels of DNA tetramers are not directly correlated with molecular size, similar to the case of natural nucleotides (19) where the blockade current was found to be determined by the competition of steric and base stacking interactions (20).

We next investigated the ability of MspA pores to resolve different tetramers containing both natural and chemically modified nucleotides (**Figure 3D**). Here, we specifically focused on heterotetramers containing a single chemically modified nucleotide (B2, B3, or B5) added in different positions of the directional sequence ACT (19). Our results clearly show that different positions of the chemically modified nucleotide in the tetramer generates distinct residual currents. For example, the residual current of heterotetrameric sequences of ACT containing four different positions of B2 (2ACT, A2CT, AC2T, and ACT2) are readily resolved at both 150 mV and 180 mV (**Figure 3D**). Although the residual current of homotetramer B2 and heterotetramer 2ACT overlap by ~29% in their Gaussians at 150 mV, they are distinguishable at 180 mV. In addition, nearly all heterotetrameric sequences of ACT containing four different positions of B3 were resolved from the homotetramer B3 at 150 and 180 mV, whereas the residual currents of 3ACT and ACT3 were only distinguishable at 180 mV (**Figure 3D**). These results are consistent with prior work reporting that tuning the applied bias is a useful approach to enhance the accuracy of nanopore-based sequencing methods (21). In summary, these results show the ability of MspA nanopores to accurately identify sequences containing chemically modified nucleotides.

In theory, sequence context allows for high-resolution readout of arbitrary combinations and arrangements of natural and modified nucleotides (A, C, G, T, B1-B7). Although specific sets of tetramers might be confused during MspA reading, the method of shift reconciliation (22) allows for such sequences to be fully resolved using the information provided by different shifts of the tetramers within the constriction of the nanopore (**Figure 3E**). The concept of shift reconciliation is illustrated with the following example, where we consider a heterogeneous sequence of 23223. In terms of the corresponding residual current levels, the prefix tetramer 2322 is confusable with 2332 or 2323 at 150 mV. However, by shifting the sliding window one position to the right, we obtain the tetramer 3223 which is not confusable with any other block. Because the trimer prefix of 3223, 322, only matches the trimer suffix of only one of the tetramers 2322, 2332, 2322 (i.e., the first one), we unambiguously deduce that 2322 is the correct prefix tetramer.

Moving beyond tetramer detection via MspA, we demonstrate that commercially available nanopore-based sequencing technology (ONT GridION) can be used to classify/sequence oligos containing the proposed molecular alphabet. For GridION experiments, the same ssDNA oligos used in MspA experiments were extended at the 3’ terminus with a polyA tail of random length >100 nts, which is used to increase the length of the oligos and guide them inside the pore (**Figure 4A**). We retrieved raw current signals from the GridION platform following a custom RNA sequencing protocol (Methods). We processed the raw current signals using deep learning techniques to discriminate and identify different combinations and orderings of the chemically modified nucleotides. As a first step, we isolated regions in the raw current signals corresponding to chemically modified nucleotides. For this purpose, we could not use the specialized software suite Tombo (23), designed by ONT for identifying potentially modified nucleotides from nanopore sequencing data, as it requires basecalling, alignment and further downstream processing. Accurate basecalling of chemically modified nucleotides is difficult to accomplish which greatly complicates alignment and classification tasks for arbitrary sub-regions of the signal. Moreover, the most recent ONT basecaller, Bonito, based on convolutional neural networks, is trained and specialized to work for natural DNA only (24). For these reasons, we developed an analysis framework that directly operates on raw current signals of the chemically modified nucleotides.

**Figure 4.**
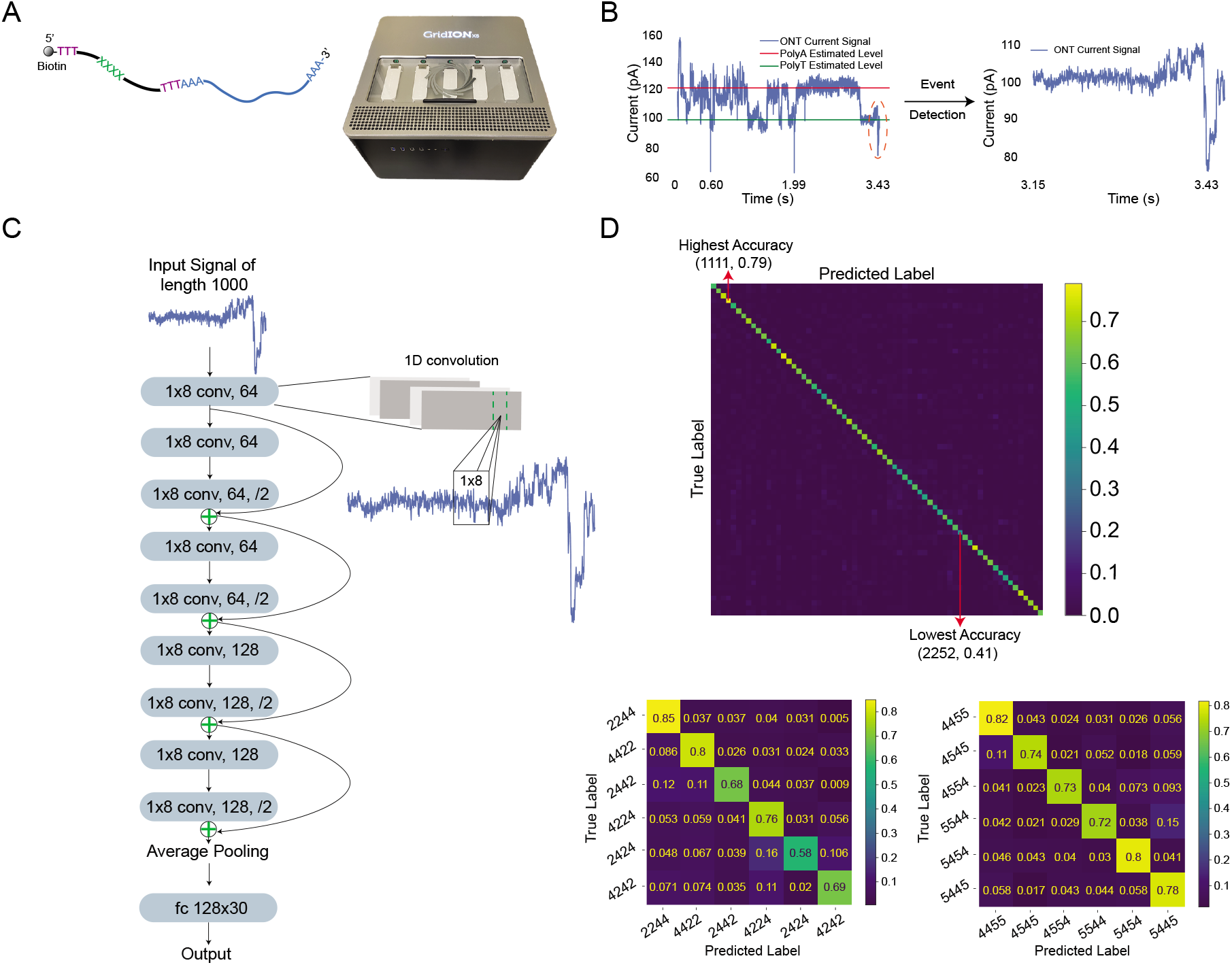
Sequencing oligos containing chemically modified nucleotides using ONT GridION. **(A)** Schematic of oligo design and a picture of the GridION sequencer used in our experiments. **(B)** (Left) Illustration of current levels of polyA and polyT regions, used in our custom level-calibration scheme. Dashed orange circle indicates the region harboring the signals from chemically modified nucleotides. (Right) Region-of-interest in raw current signal obtained by identifying polyA-polyT patterns. **(C)** Neural network model used for classification. The 1D residual neural network architecture comprises nine 1D convolution blocks. For example, a 1D convolution block (1×8 conv, 64) indicates that the kernel size for the convolution is 1×8 and that the number of output channels is 64. Half-downsampling for each channel is denoted by (/2); averaging over all channels to arrive at a single vector is referred to as “Average Pooling”; the (fc 128×30) notation indicates a fully connected layer with the shape 128×30. (Right) Magnified view of the operation of 1D convolutional neural networks on time-series data. **(D)** (Top) Confusion matrix for 66 classes, all of which have roughly the same number of samples (subsampled to ~3500 sample oligos in each class). Random guessing would lead to a classification accuracy of 1.52%, whereas the smallest accuracy from our model is 41% (tetramer 2252). For our model-based prediction, the mean classification accuracy is 60.28% ± 0.28% (39X larger than random guessing), and the highest observed accuracy is 79% (tetramer 1111). The exact number of samples in each class is listed in **Table S5**. (Bottom left) Confusion matrix for six selected classes using B2 and B4 (named as listed, subsampled to roughly 5000 samples per class). Random guessing leads to an accuracy of 16.67%, whereas our modelbased prediction ensures an average classification accuracy of 72.25% ± 1.46%. (Bottom right) Confusion matrix for six selected classes using B4 and B5 (named as listed, subsampled to roughly 5000 samples per class). Random guessing leads to an accuracy of 16.67%, while our model-based prediction ensures an average accuracy of 77.84% ± 0.96%.

Analysis of raw current signals is challenging because nanopore current signals exhibit extreme variations known as level drifts (**Figure S5**). Level drifts arise because each membrane patch (recording channel) inside the device has its own electric circuit, and each pore has unique features. To address this challenge, we developed a two-step identification scheme depicted in **Figure 4B**. In the first step, we estimate the current level for the polyA region, and subsequently use it for signal calibration. Similar calibration steps are standardly performed for nanopore sequencing of natural DNA, but they rely on adaptor-based calibrations since all analytes use identical adaptors with a well-defined sequence content. For actual level calibration, we used kernel density estimation of the signal level distribution (25), followed by identification of the levels that have the two largest probabilities in the estimated distribution. This approach is justified because polyA regions constitute the longest signal component in our oligo sequences. Moreover, on average, polyT levels are expected to be lower than polyA levels, so readout regions that are trailed by nearly flat regions with a mean level value lower than that for the polyA tails are filtered using a finite state machine (26). These regions are expected to bear signals from the chemically modified nucleotides. After extracting modification-bearing signals, raw current readouts are subsequently classified. For this task, we designed a 1D residual neural network model (27,28) (**Figure 4C**) containing 1D convolution layers (conv) that serve as feature extractors, and one fully connected layer (fc) that serves as a classifier. The model is trained on oligo data corresponding to different combinations and orderings of chemically modified nucleotides, with each option supported by thousands of training samples (**Table S5**). Elements from each class are uniformly sampled at random in a balanced manner and split into training/validation/test sets with splitting percentages 60%/20%/20%, respectively.

Results from neural network-guided identification tasks pertaining to five independent experimental runs are shown in **Figure 4D**. Confusion matrices are used to summarize the prediction accuracies, ranging between 0 and 1 (with 1 corresponding to perfectly accurate identification). Importantly, these results show that most tetramers are identified with high accuracy (i.e., the diagonal elements are significantly larger than the off-diagonal elements). The average classification accuracy for each model is provided in the caption of **Figure 4D**, along with the accuracy one would expect from random guessing. For example, we observed an accuracy of 0.85 for heterotetramers (2244, 2244), which is to be interpreted as an 85% success rate in correctly identifying the sequence 2244, or a 15% chance of misinterpreting 2244 as another combination or sequence order (**Figure 4D**). Overall, we performed a total of 13 different classification tasks, including one task for all classes (77 in total, from which only 66 were depicted due to small amounts of training data for the remaining 11 classes). We further included 12 tasks involving subsets of classes containing chemically modified nucleotides shown in **Figure 1**. For brevity, two results for 2X+2Y classes and a summary of all results are shown in **Figure 4D**; the full set of results are shown in **Figure S6**.

In closing, we report an expanded alphabet for DNA data storage compatible with nanopore sequencing technology. The unique feature of our approach is coupled, iterative selection and testing that involves determining suitability for forming stable duplex structures and nanopore sequencing. Overall, the described system enables the recording of digital data with increased storage density and more bits per synthesis cycle. In particular, our storage system enables a maximum recording density of log_2_11 bits in each cycle, compared to log_2_ 4 = 2 bits for natural DNA; this strategy also theoretically increases the rate (speed) of the recorder by 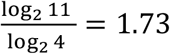 fold. Our extensive nanopore experiments provide strong evidence that many more chemically modified nucleotides can be used for molecular storage because many ionic current levels remain available, i.e., the ionic current spectrum is sparsely populated. In addition, our system allows for high-fidelity readouts and PCR-based random-access features for encodings restricted to duplex formation competent monomers. Although not all pairings of chemical modifications may be suitable for amplification using natural enzymes, and some duplex formations may be unstable, the proposed system provides the first example of a coupled coding alphabet and channel selection and optimization paradigm. In conclusion, this work demonstrates fundamentally new directions in molecular storage that hold the potential to advance the field of DNA-based data storage.

## Acknowledgements

The work was funded by the NSF+SRC SemiSynBio program under agreement number 1807526 and NSF grants 1618366 and 2008125. A.A. and M.C. acknowledge support from NHGRI/NIH via grant R21-HG011741. The supercomputer time was provided by the University of Illinois at the Blue Waters Petascale System. The authors gratefully acknowledge many useful discussions with Profs. Jean-Pierre Leburton and Xiuling Li, as well as Nagendra Athreya and Apratim Khadelwal and Dr. Bo Li.

## Author Contributions

O.M. and C.M.S. conceived the ideas for expanded DNA alphabets. S.K.T., O.M., C.M.S., M.C., and B.P. designed the ssDNA oligos containing the modified nucleotides. M.C., B.P. and S.A.S. designed and performed the MspA nanopore experiments. A.G.H. designed and performed the ONT sequencing experiments. C.P., O.M., S.C. and A.G.H performed the ONT raw output data analysis. J.L. and A.A. designed and performed the MD simulations. All authors contributed towards the system development and participated in the writing of the manuscript.

## Materials and Methods

### Oligo design and synthesis

All oligos tested are of fixed length 40nt and synthesized by Integrated DNA Technologies (IDT). For MspA experiments, the content of the oligos was chosen to include two polyT sequences at locations 1-12 and 17-40, and a chemically modified tetramer at positions 13-16. All oligos were biotinylated at the 5’ end.

### PCR Amplification

DNA amplification was performed via PCR using Q5 DNA polymerase, 5× Q5 buffer and pUC19 plasmid as template (New England Biolabs) in 50 μl. The 1.4kb sequence is: 5’CGTTTTACAACGTCGTGACTGGGAAAACCCTGGCGTTACCCAACTTAATCGCCTTGCAGCACATCCCCCTTTCGCCAGCTGGCGTAATAGCGAAGAGGCCCGCACCGATCGCCCTTCCCAACAGTTGCGCAGCCTGAATGGCGAATGGCGCCTGATGCGGTATTTTCTCCTTACGCATCTGTGCGGTATTTCACACCGCATATGGTGCACTCTCAGTACAATCTGCTCTGATGCCGCATAGTTAAGCCAGCCCCGACACCCGCCAACACCCGCTGACGCGCCCTGACGGGCTTGTCTGCTCCCGGCATCCGCTTACAGACAAGCTGTGACCGTCTCCGGGAGCTGCATGTGTCAGAGGTTTTCACCGTCATCACCGAAACGCGCGAGACGAAAGGGCCTCGTGATACGCCTATTTTTATAGGTTAATGTCATGATAATAATGGTTTCTTAGACGTCAGGTGGCACTTTTCGGGGAAATGTGCGCGGAACCCCTATTTGTTTATTTTTCTAAATACATTCAAATATGTATCCGCTCATGAGACAATAACCCTGATAAATGCTTCAATAATATTGAAAAAGGAAGAGTATGAGTATTCAACATTTCCGTGTCGCCCTTATTCCCTTTTTTGCGGCATTTTGCCTTCCTGTTTTTGCTCACCCAGAAACGCTGGTGAAAGTAAAAGATGCTGAAGATCAGTTGGGTGCACGAGTGGGTTACATCGAACTGGATCTCAACAGCGGTAAGATCCTTGAGAGTTTTCGCCCCGAAGAACGTTTTCCAATGATGAGCACTTTTAAAGTTCTGCTATGTGGCGCGGTATTATCCCGTATTGACGCCGGGCAAGAGCAACTCGGTCGCCGCATACACTATTCTCAGAATGACTTGGTTGAGTACTCACCAGTCACAGAAAAGCATCTTACGGATGGCATGACAGTAAGAGAATTATGCAGTGCTGCCATAACCATGAGTGATAACACTGCGGCCAACTTACTTCTGACAACGATCGGAGGACCGAAGGAGCTAACCGCTTTTTTGCACAACATGGGGGATCATGTAACTCGCCTTGATCGTTGGGAACCGGAGCTGAATGAAGCCATACCAAACGACGAGCGTGACACCACGATGCCTGTAGCAATGGCAACAACGTTGCGCAAACTATTAACTGGCGAACTACTTACTCTAGCTTCCCGGCAACAATTAATAGACTGGATGGAGGCGGATAAAGTTGCAGGACCACTTCTGCGCTCGGCCCTTCCGGCTGGCTGGTTTATTGCTGATAAATCTGGAGCCGGTGAGCGTGGGTCTCGCGGTATCATTGCAGCACTGGGGCCAGATGGTAAGCCCTCCCGTATCGTAGTTATCTACACGACGGGGAGTCAGGCAACTATGGATGAACGAAATAGACAGATCGCTGAGATAGGTGCCTCACTGATTAAGCATTGGTA3’.

All primers were purchased from Integrated DNA Technologies (IDT). Both B1 and B2 were purchased from TriLink Biotechnologies in form of triphosphates (https://www.trilinkbiotech.com/2-amino-2-deoxyadenosine-5-triphosphate-n-2003.html and https://www.trilinkbiotech.com/5-hydroxymethyl-2-deoxycytidine-5-triphosphate.html). All natural and chemically modified nucleotides were added in equimolar ratios in all PCR reactions.

### MD Simulations

The molecular mechanics models of modified nucleotides B1, B3, B4, B5 and B6, including their topology and force field parameter files, were generated using the CHARMM General Force Field (CGenFF) (30). The charge of the atom connecting to the sugar was adjusted so that the total charge of the base is zero, which is the case for all the natural nucleotides in CHARMM36. The parameters for B2 were adopted from a previous study (31). Eight systems each containing a modified Dickerson dodecamers (CGCGAATTCGCG)(15) were created starting from a B-DNA conformation to contain two different pairs of modified and natural bases while all other bases remained as in the original sequence. Each DNA duplex was immersed in a 75 Å x 75 Å x 75 Å volume of 1M KCl solution. After 2000 steps of energy minimization, the systems were equilibrated with the DNA backbone phosphate atoms restrained (*k_s_* = lkcal/mol/Å^2^) for the first 10ns. Each system contains approximately 39,000 atoms. Additional restrains were applied to enforce the expected hydrogen bonds between the modified and natural nucleotides for the first 20 ns. The systems were simulated for 350 ns in the absence of any restrains in the constant number of particles, pressure (1 atm) and temperature (295 K) ensemble using NAMD2 (32). If prominent structural disruptions had developed in both base pairs surrounding the modified nucleotide base pair, the simulation was terminated. Specifically, the simulation of the systems containing the B4 nucleotide lasted only 250 ns. Simulations of all the systems were performed using periodic boundary conditions. The simulations employed the particle mesh Ewald (PME) algorithm (33) to calculate long-range electrostatic interaction over a 1Å-spaced grid. RATTLE (34) and SETTLE (35) algorithms were adopted to constrain all covalent bonds involving hydrogen atoms, allowing 2-fs time step integration used in the simulations. van der Waals interactions were calculated using a smooth 10 – 12 Å cutoff. The NPT ensembles used the Nosé-Hoover Langevin piston pressure control (36), which maintained a constant pressure by adjusting system’s dimension. Simultaneously, Langevin thermostat (25) was adopted for temperature control, with damping coefficient of 0.5 ps^_1^ applied to all heavy atoms in the systems. CHARMM36 (37), output of CGenFF (30), TIP3P water model (38) as long as custom NBFIX corrections to nonbonded interactions (39) were employed as the parameter set of the simulation. The hydrogen bonds occupancy, the distances between hydrogen bond donors and acceptors as well as the short/long axis lengths of bases are calculated from the well equilibrated last 100 ns fragment of the trajectory using VMD (40). The hydrogen bonds were defined to have the donor-accepter interaction distance of less than 3Å and the cutoff angle of 20°. Given the largely planar shape of the bases, their short/long were determined by first computing the three principal axes of the bases and then choosing the largest two values. Simulations/analysis of the B4 pairing with natural bases in longer DNA strands were conducted using the same methodology, but with only one modified base contained in the dodecamer. Besides, extra bonds were applied to the donor(N1) and accepter(N3) atoms on the terminal pairs to prevent the ends from fraying in these simulations to adapt the situation of long DNA strands. These simulations ran 550ns except if unstable configurations were observed.

### MspA nanopores and purification of M2-NNN MspA

All chemicals were purchased from Fisher Scientific unless stated otherwise. Streptavidin was ordered from EMD Millipore (Burlington, MA) (Catalog # 189730). Phenylmethylsulfonyl fluoride (PMSF) was ordered from GoldBio (St. Louis, MO) (Catalog # P-470). DNA of M2-NNN MspA construct(29) was a gift from Dr. Giovanni Maglia (University of Groningen, Netherlands). The pT7-M2-NNN-MspA was transformed into BL21 (DE3) pLyss cells and grown in LB medium at 37°C until the OD600 reached 0.5-0.6. The cells were then induced with 0.5 mM isopropyl β-D-1-thiogalactopyranoside (IPTG) and continued to grow at 16°C for 16 hours. Cells were harvested and centrifuged at 19,000 x g for 30 min at 4°C. Cells were resuspended in the lysis buffer containing 100 mM Na_2_HPO_4_/NaH_2_PO_4_, 1 mM ethylenediaminetetraacetic acid (EDTA), 150 mM NaCl, 1 mM phenylmethylsulfonyl fluoride (PMSF) pH 6.5, before heating at 60°C for 10 minutes. The cells were sonicated by using VWR Scientific Branson 450 sonicator (duty cycle of 20% and output control of 2) for 8 minutes. The lysate was centrifuged at 19,000 x g for 30 min and the supernatant was discarded. The pellet was resuspended in the solubilization buffer containing 100 mM Na_2_HPO_4_/NaH_2_PO_4_, 1 mM EDTA, 150 mM NaCl, 0.5% (v/v) Genapol X – 80, pH 6.5. After completely resuspending the pellet, it was centrifuged at 19,000 x g for 30 min. The supernatant, containing solubilized membrane extract, was collected for Ni-NTA purification. MspA was further purified using a 5 mL HisPur™ Ni-NTA resin (GE Healthcare) and eluted in a buffer of 0.5 M NaCl, 20 mM HEPES, 0.5% (v/v) Genapol X – 80, pH 8.0 by applying an imidazole gradient. MspA oligomers were further purified by SDS-PAGE gel extraction. The purified MspA protein was run in 7.5% SDS-PAGE gel. The band of MspA oligomer was cut from the gel and extracted in the extraction buffer containing 50 mM Tris-HCl, 150 mM NaCl, 0.5% Genapol X – 80, pH 7.5. The protein was extracted at room temperature (23°C) for 6 hours before centrifuged at 9,000 x g for 30 min to collect the protein solution. The purified MspA oligomer was fast frozen and stored at −80°C for further use.

### Single-channel recording using MspA

The experiments were performed in a device containing two chambers separated by a 25 μm thick polytetrafluoroethylene film (Goodfellow) with an aperture of approximately 100 μm diameter located at the center. A hexadecane/pentane (10% v/v) solution was first added to cover both sides of the aperture. After the pentane evaporated, each chamber was then filled with buffer containing 1 M KCl 10 mM HEPES pH 8.0. 1,2-diphytanoyl-sn-glycero-3-phosphocholine (DPhPC) dissolved in pentane (10 mg/mL) was dropped on the surface of the buffer in both chambers. After the pentane evaporated, the lipid bilayer was formed by pipetting the solution in both chambers below the aperture several times. An Ag/AgCl electrode was immersed in each chamber with the *cis* side grounded. M2-NNN MspA proteins (around 1 nM, final concentration) were also added to the *cis* chamber. To promote MspA insertion, a ? +200 mV voltage was applied. After a single MspA was inserted into the planar lipid bilayer, the applied voltage was decreased to 150 mV (or 180 mV) for recording. The current was amplified with an Axopatch 200B integrating patch-clamp amplifier (Axon Instruments, Foster City, CA). Signals were filtered with a Bessel filter at 2 kHz and then acquired by a computer (sampling at 100 μs) after digitization with a Digidata 1440A/D board (Axon Instruments).

### DNA immobilized in MspA

After recording a single MspA pore for 5-10 minutes at positive voltages to check its stability, 5’-biotinylated DNA sample (final concentration of 0.25 μM) was added to the *cis* chamber. Streptavidin (0.1 μM), added to solutions in the *cis* chamber, can bind to biotin to prevent the full translocation of the DNA strand through the nanopore. To collect the signal generated from each DNA samples, we applied a sweep protocol. The amplifier applied either 150 mV or 180 mV for 10 s then applied −150 mV to force the DNA out of the pore back into the *cis* compartment. The voltage was then returned to the original value and the sweep protocol repeated for at least 40 times at each voltage.

### ONT sequencing protocol

NEB terminal transferase was used for A-tailing the 3’ end of the 40-mer control oligos. The reaction mixture was made by 5ul 10X TdT buffer, 5ul 2.5mM CoCl2, 5 pmole DNA, 0.5ul 10mM dATP, 0.5 ul terminal transferase, and 38 ul H_2_O. The reaction was Incubated at 37 C for 30 mins, followed by inactivation at 70 C for 10 mins. The DNA was then purified using the Zymo DNA clean up kit (ssDNA Buffer:sample=7:1) and eluted in 10ul warm H_2_O. The Oxford Nanopore SQK-RNA002 kit was used for library preparation. The RT adaptor was ligated for 10min at room temperature, then mixed with reverse transcription master mix. 2uL of Superscript IV were added and the mixture was Incubated at 50 C for 50mins, followed by 70 C for 10mins and cooled down to 4 C. Bead clean-up was performed using 40ul samples with 72ul RNAClean XP beads, rotated for 5mins, washed by 70% EtOH and eluted by 20ul H_2_O. The RMX adaptor was ligated in 10mins at room temperature, then 40ul RNA Clean XP beads clean-up was used, and the product was washed with 150ul of the wash buffer twice. It was then eluted in 21ul of the elution buffer. The reaction was loaded onto an R9.4.1 flowcell and sequenced on a GridION X5 (Oxford Nanopore) for 24 hs.

