## Supplementary Information for "Expanding the Molecular Alphabet of DNA-Based Data Storage Systems with Neural Network Nanopore Readout Processing"

*Tabatabaei et al.*

**This file contains:**

Supplementary Notes

Figures S1-S6

Tables S1-S5

References

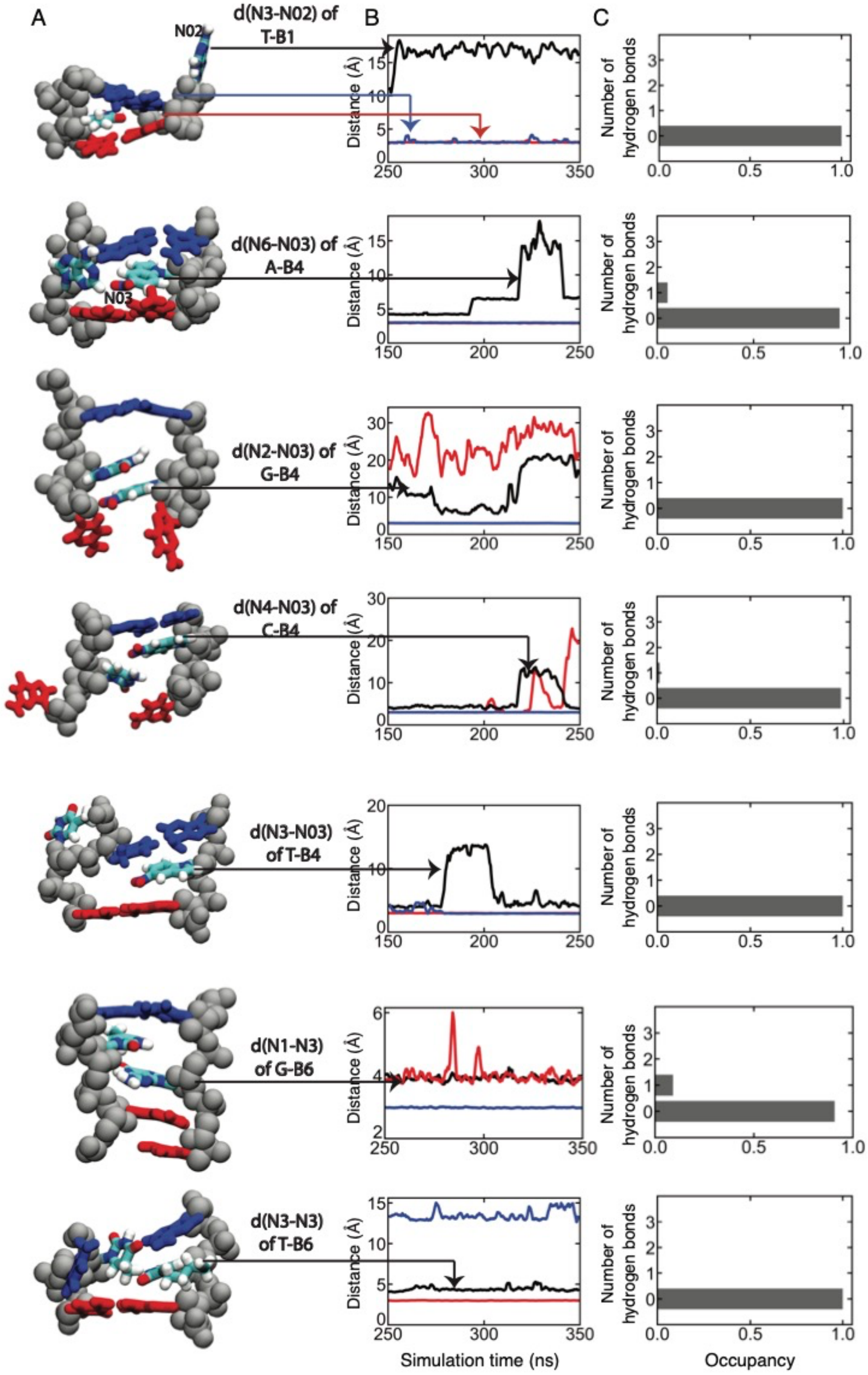
A
B
C
N02
d(N3-N02) of T-B1
N03
d(N6-N03) of A-B4
d(N2-N03) of G-B4
d(N4-N03) of C-B4
d(N3-N03) of T-B4
d(N1-N3) of G-B6
d(N3-N3) of T-B6
Distance (Å)
Simulation time (ns)
Number of hydrogen bonds
Occupancy

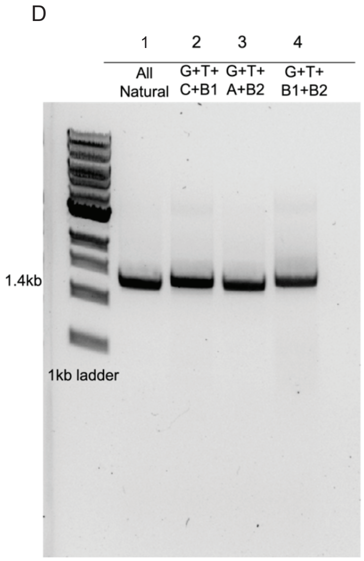


**Figure S1**. Interactions between modified and natural bases that do not involve stable hydrogen bonds. **(A)** Microscopic configurations of modified base pairs (from top to bottom: B1—T, B2—G, A—B4, G—B4, C—B4, T—B4, G—B6, and T—B6). The backbone of the dodecamer is shown using silver spheres whereas the bases are drawn as molecular bonds. Unnatural bases and the natural bases that pair with them are colored according to the atom type (cyan for carbon, blue for nitrogen and red for oxygen). Base pairs immediately adjacent to the modified base pair are colored in red or blue.

**(B)** Distance between the key atoms of the modified base pair during the last 100 ns of the 350 ns MD simulation. The red curve and blue curve show the N1—N3 distance for the two adjacent base pairs, whose pairing patterns can either remain intact or be disrupted. The arrows starting from panel A to panel B indicate the correspondence between the basepairs and the curves. The label specifies the atoms used to compute the distance. The curves show a running average of the 10 ps-sampled data with a 2 ns averaging window. **(C)** Probability of observing the specified number of hydrogen bonds within a modified base pair. The H-bonding probabilities were computed using the final 100 ns of a 350 ns all-atom MD simulation of a DNA dodecamer. **(D)** As a starting point to experimentally evaluate the effect of chemically modified nucleotides on DNA structure, we performed a simple PCR reaction on a 1.4kb double stranded DNA from a commonly used vector, pUC19 plasmid, using Q5 polymerase. The reaction was either supplied by all four natural nucleotides or B1 and B2 as substitutes for A and C. The final PCR products were run on 1% agarose gel. The results indicate successful incorporation of B1 and B2 into DNA duplex structure when only one of them (lanes 2 and 3) or two of them (lane 4) were used instead of the natural nucleotides.

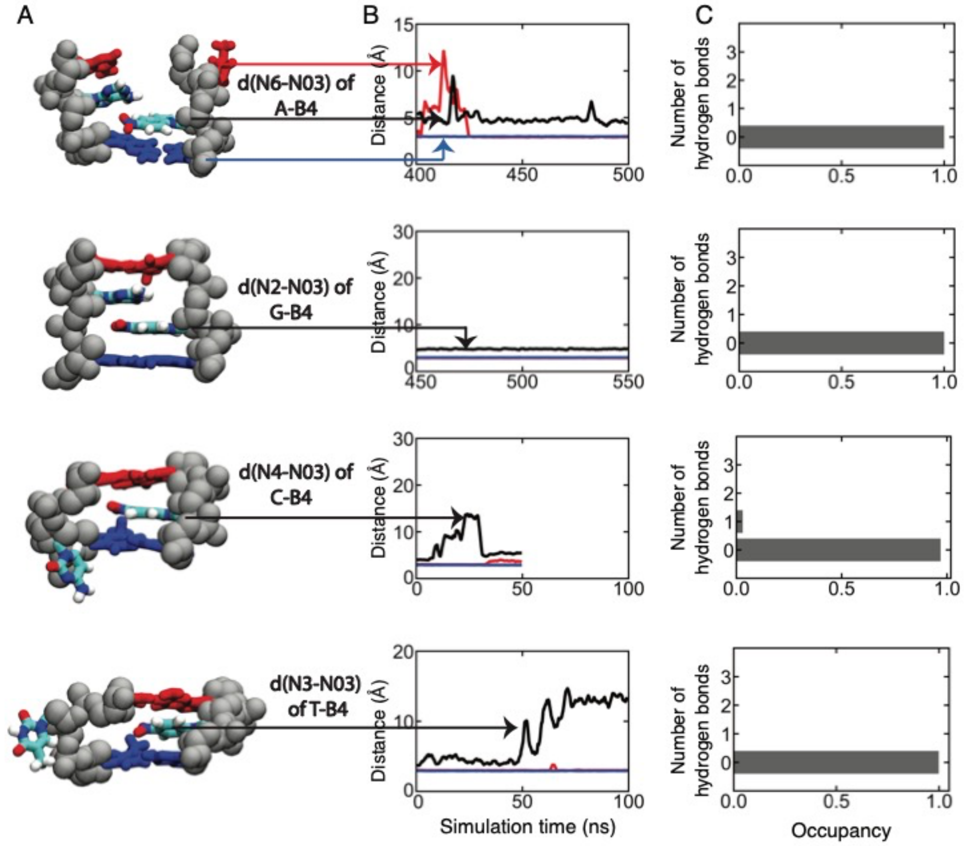


**Figure S2**. Interactions between B4 and natural bases in long DNA strands **(A)** Microscopic configurations of modified base pairs (from top to bottom: A—B4, G—B4, C—B4 and T—B4). The backbone of the dodecamer is shown using silver spheres whereas the bases are drawn as molecular bonds. B4 bases and the natural bases that pair with them are colored according to the atom type (cyan for carbon, blue for nitrogen and red for oxygen). Base pairs immediately adjacent to the modified base pair are colored in red or blue. In contrast to simulations reported in Figure S4, here each DNA dodecamer contains contains only one B4 base. Extra bonds between donor(N1) and acceptor(N3) (The equilibrium length was set as 2.9 Å. The spring constant was set as $1 \mathrm{kcal/mol/Å^2}$.) are applied the terminal base pairs, preventing DNA from fraying and thereby mimicking an environment of a longer DNA strand. **(B)** Distance between the key atoms of the modified base pair during the last 50/100 ns of the MD simulation. The red curve and blue curve show the N1—N3 distance for the two adjacent base pairs, whose pairing patterns can either remain intact or be disrupted. The arrows starting from panel A to panel B indicate the correspondence between the basepairs and the curves. The label specifies the atoms used to compute the distance. The curves show a running average of the 10 ps-sampled data with a 2 ns averaging window. **(C)** Probability of observing the specified number of hydrogen bonds within a modified base pair. The H-bonding probabilities were computed using the final 50/100ns of the all-atom MD simulations of a DNA dodecamer.

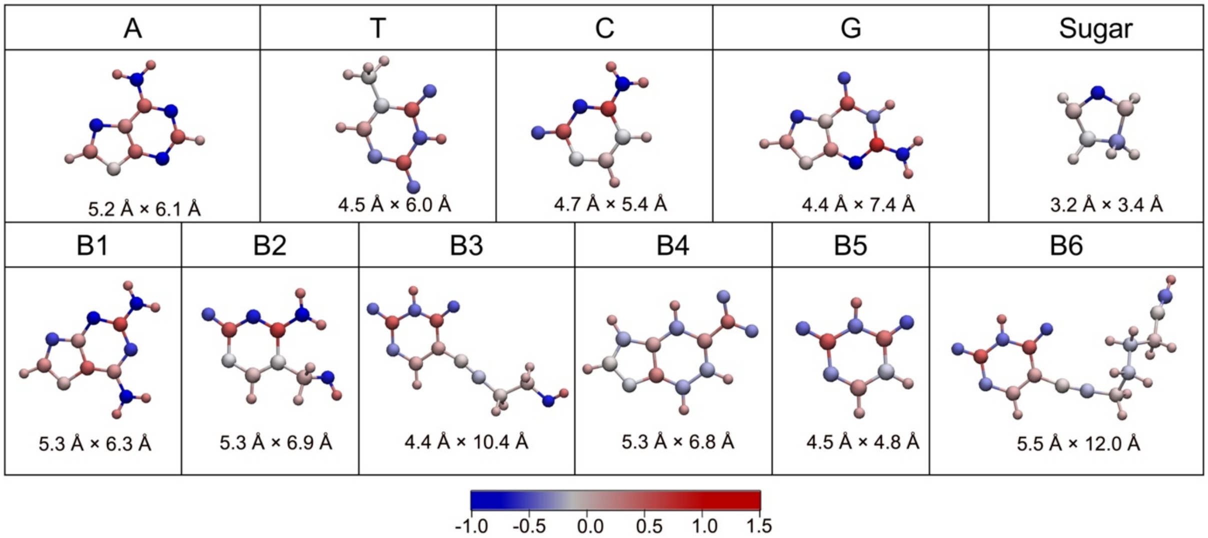


**Table S1**. Charge distribution and dimensions of the natural and modified bases and deoxyribose moieties in simulation. The chemical moieties are shown using a ball-and-stick representation, with the atoms colored by their charge according to the color bar. The dimensions of each base, specified as short axis length × long axis length, were averaged over the last 100 ns of a 350 ns/250ns all-atom MD trajectory of a DNA dodecamer.

Although it may appear that issues observed with some monomers may prohibit the use of them in storage applications, or implementations that include the other recommended chemically modified nucleotides, further experiments are required to confirm which combinations may cause disruptions. Once such combinations are identified, well-known methods from coding theory – such as constrained coding (1) – may be used to eliminate the offending patterns with minimal loss in the information rate.

### Further results on the MspA readout experiments

Here, we provide a complete report on the experimental results of MspA nanopore detection of all 77 chemically modified tetramers.

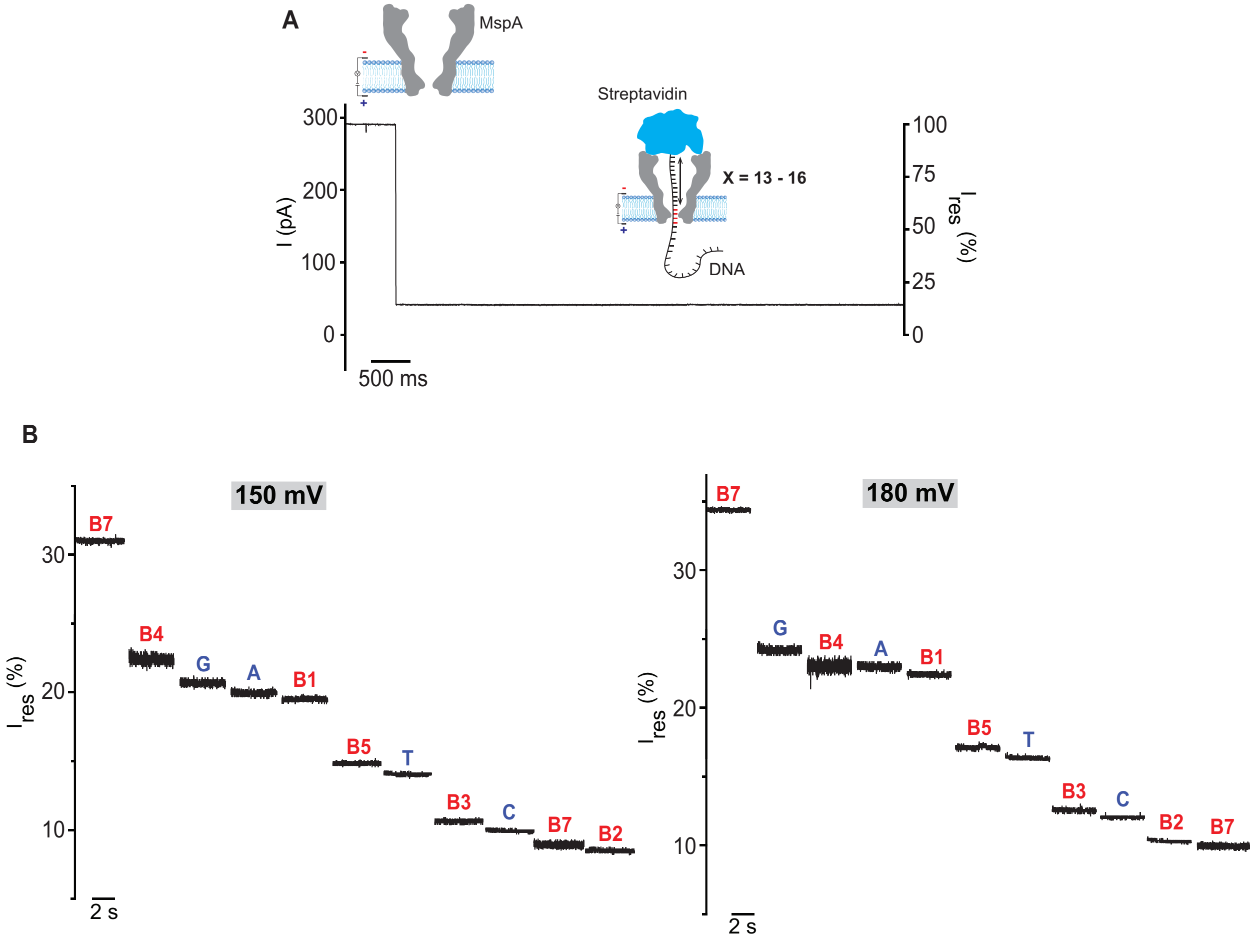


**Figure S3. Discrimination of immobilized DNA by MspA nanopore. (A)** Schematic diagram of DNA immobilized in the MspA nanopore. Single-stranded DNA (ssDNA) was attached to a streptavidin molecule (cyan) using a biotin linker. Bulky streptavidin prevents ssDNA to translocate through the MspA pore (gray). The residual ion current was recorded as the ssDNA is immobilized within the pore, which is generated by 4 nucleotides in and around the constriction side, at positions 13 – 16 from the biotin-streptavidin end. The open-pore current of MspA is normalized to 100%. **(B)** The representative single-channel recording generated by each tetramer sequence at positions 13-16 from the tethering point to the constriction site (reading head) of the MspA pore. Native nucleotides are highlighted in blue and modified nucleotides in red. Buffer used is 1 M KCl 10 mM HEPES pH 8.0.

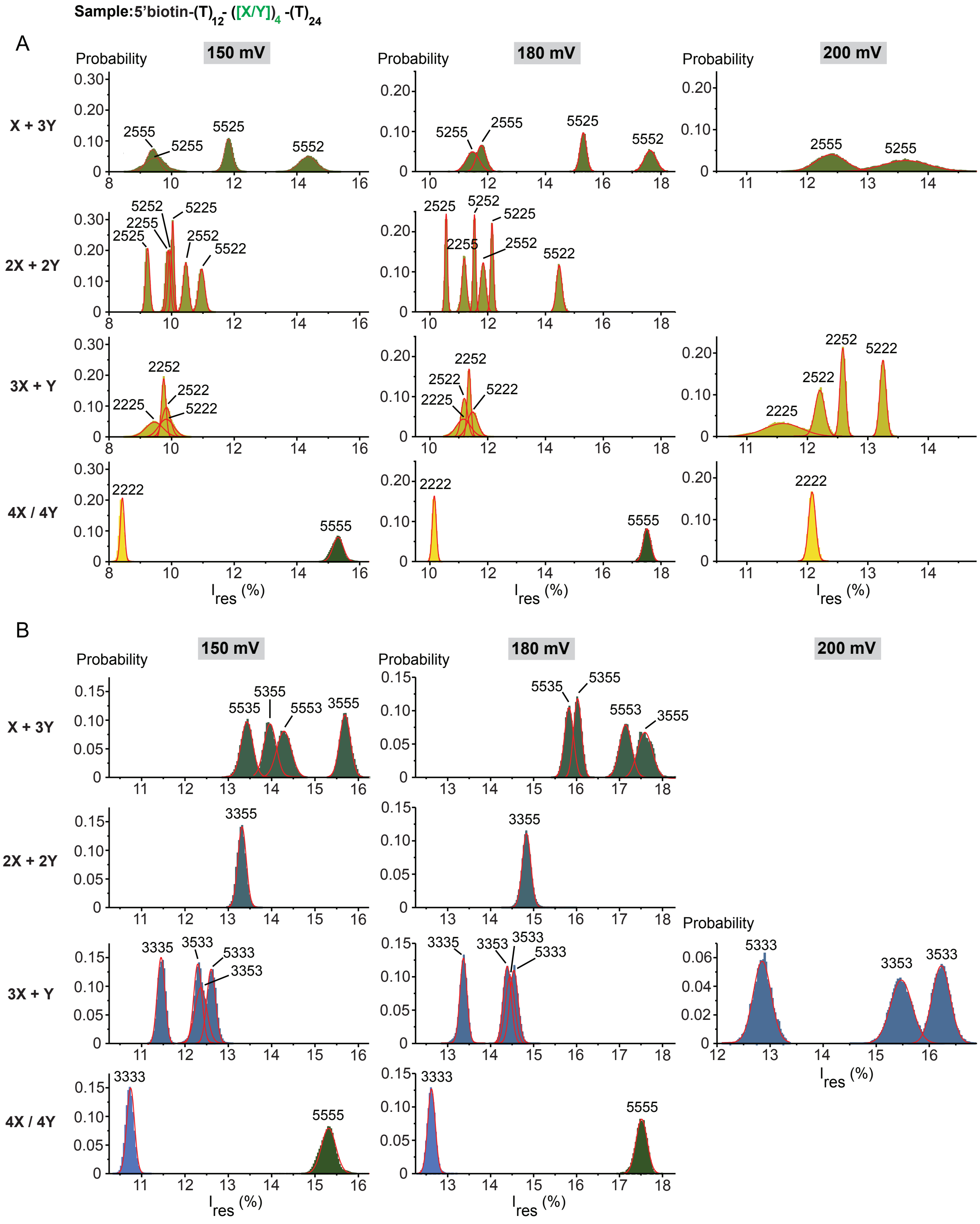
Sample:5'biotin-(T)12-([X/Y])4-(T)24
A
B
150 mV
180 mV
200 mV
Probability
X + 3Y
2X + 2Y
3X + Y
4X / 4Y
Ires (%)
2555
5255
5525
5552
2525
2255
5252
5225
2552
5522
2225
2252
2522
5222
2222
5555
5535
5355
5553
3555
3355
3335
3533
5333
3353
3333

C

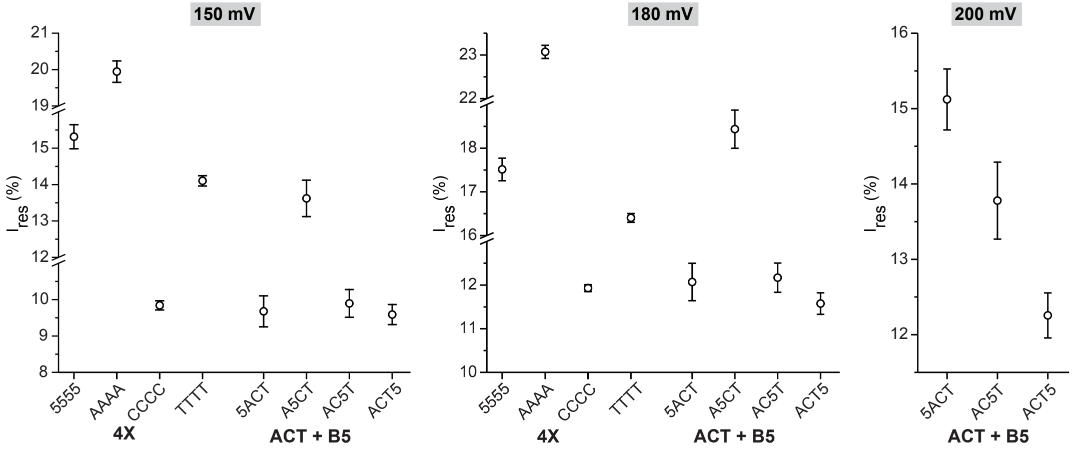


**Figure S4.** Histograms of the averaged residual ionic currents and the fitted Gaussian curves at various applied voltages for tetramers involving different orderings of B2 and B5 monomers **(A)** and B3 and B5 monomers **(B)** at 150, 180, and 200 mV. All experiments were performed in aqueous buffer (1 M KCl 10 mM HEPES pH 8.0).

**(C)** Peak values and full-width half-height values (FWHM), represented as error bars, of the fitted Gaussian distributions around mean residual ionic currents generated by different orderings of B5 with the natural nucleotides (A, C, and T) at 150,180, and 200 mV. All experiments were performed in aqueous buffer (1 M KCl 10 mM HEPES pH 8.0).

**Table S2.** The mean residual currents ($I_{res}$ (%)) and the full-width half-height (FWHM) values for each oligonucleotide were determined by Gaussian fitting of the residual current histogram from experiments with different combination of natural and modified nucleotides at positions 13 – 16 from the streptavidin anchor at 150 mV.

| Combination | X | Y | Sample | $I_{res}$ (%) | FWHM |
|---|---|---|---|---|---|
| ACT+X | 2 | | 2ACT | 8.68 | 0.60 |
| | | | A2CT | 10.65 | 0.22 |
| | | | AC2T | 10.14 | 0.67 |
| | | | ACT2 | 9.01 | 0.32 |
| | 3 | | 3ACT | 9.60 | 0.36 |
| | | | A3CT | 10.27 | 0.70 |
| | | | AC3T | 8.69 | 0.41 |
| | | | ACT3 | 9.52 | 0.48 |
| | 5 | | 5ACT | 9.68 | 0.43 |
| | | | A5CT | 13.62 | 0.50 |
| | | | AC5T | 9.90 | 0.38 |
| | | | ACT5 | 9.59 | 0.28 |
| 4X | 1 | | B1 | 19.66 | 0.39 |
| | 2 | | B2 | 8.43 | 0.13 |
| | 3 | | B3 | 10.75 | 0.18 |
| | 4 | | B4 | 22.74 | 0.51 |
| | 5 | | B5 | 15.32 | 0.33 |
| | 6 | | B6 | 8.49 | 0.29 |
| | 7 | | B7 | 31.30 | 0.12 |
| | A | | A | 19.94 | 0.29 |
| | C | | C | 9.84 | 0.13 |
| | G | | G | 20.82 | 0.50 |
| | T | | T | 14.10 | 0.14 |
| 3X+Y | 2 | 3 | 2223 | 9.13 | 0.34 |
| | | | 2232 | 7.36 | 0.38 |
| | | | 2322 | 8.34 | 0.37 |
| | | | 3222 | 9.45 | 0.29 |
| | | 5 | 2225 | 9.45 | 0.55 |
| | | | 2252 | 9.75 | 0.14 |
| | | | 2522 | 9.83 | 0.27 |

| | | | | | |
|---|---|---|---|---|---|
| | | | 5222 | 9.83 | 0.48 |
| | 3 | 2 | 2333 | 7.91 | 0.19 |
| | | | 3233 | 7.48 | 0.30 |
| | | | 3323 | 9.44 | 0.29 |
| | | | 3332 | 10.45 | 0.42 |
| | | 5 | 3335 | 11.45 | 0.18 |
| | | | 3353 | 12.37 | 0.27 |
| | | | 3533 | 12.30 | 0.19 |
| | | | 5333 | 12.61 | 0.20 |
| | 5 | 2 | 2555 | 9.39 | 0.37 |
| | | | 5255 | 9.46 | 0.60 |
| | | | 5525 | 11.80 | 0.25 |
| | | | 5552 | 14.35 | 0.55 |
| | | 3 | 3555 | 15.69 | 0.25 |
| | | | 5355 | 13.96 | 0.28 |
| | | | 5535 | 13.43 | 0.27 |
| | | | 5553 | 14.29 | 0.34 |
| **2X+2Y** | 2 | 3 | 2323 | 8.53 | 0.17 |
| | | | 2332 | 8.07 | 0.14 |
| | | | 3223 | 10.02 | 0.16 |
| | | | 3232 | 8.18 | 0.16 |
| | | | 3322 | 11.34 | 0.17 |
| | | | 2233 | 7.59 | 0.14 |
| | | 4 | 2424 | 12.79 | 0.20 |
| | | | 2442 | 13.01 | 0.59 |
| | | | 4224 | 12.39 | 0.12 |
| | | | 4242 | 12.62 | 0.19 |
| | | | 4422 | 12.99 | 0.21 |
| | | | 2244 | 10.78 | 0.18 |
| | 2 | 5 | 2525 | 9.23 | 0.13 |
| | | | 2552 | 10.45 | 0.17 |
| | | | 5225 | 10.03 | 0.09 |
| | | | 5252 | 9.95 | 0.14 |
| | | | 5522 | 10.96 | 0.20 |
| | | | 2255 | 9.89 | 0.13 |
| | 4 | 5 | 4545 | 23.07 | 0.34 |

| | | | | | |
|---|---|---|---|---|---|
| | | | 5454 | 20.16 | 0.43 |
| | | | 4554 | 19.55 | 0.20 |
| | | | 5445 | 19.38 | 0.32 |
| | | | 5544 | 17.63 | 0.24 |
| | | | 4455 | 22.01 | 0.33 |
| | 1 | 2 | 1122 | 11.18 | 0.27 |
| | | 3 | 1133 | 16.16 | 0.22 |
| | | 4 | 1144 | 18.09 | 0.30 |
| | | 5 | 1155 | 17.57 | 0.21 |
| | 3 | 4 | 3344 | 19.07 | 0.85 |
| | | 5 | 3355 | 13.32 | 0.19 |

**Table S3**. The mean residual currents ($I_{res}$ (%)) and the full-width half-height (FWHM) values for each oligonucleotide, determined by performing Gaussian fitting of the residual current histogram from experiments involving different combination of natural and modified nucleotides at positions 13 – 16 from the streptavidin anchor at 180 mV.

| Combination | X | Y | Sample | $I_{res}$ (%) | FWHM |
|---|---|---|---|---|---|
| ACT+X | 2 | | 2ACT | 11.06 | 0.49 |
| | | | A2CT | 12.93 | 0.21 |
| | | | AC2T | 12.02 | 0.59 |
| | | | ACT2 | 10.53 | 0.28 |
| | 3 | | 3ACT | 12.38 | 0.38 |
| | | | A3CT | 14.27 | 0.61 |
| | | | AC3T | 10.74 | 0.40 |
| | | | ACT3 | 11.38 | 0.42 |
| | 5 | | 5ACT | 12.07 | 0.43 |
| | | | A5CT | 18.44 | 0.44 |
| | | | AC5T | 12.17 | 0.34 |
| | | | ACT5 | 11.58 | 0.25 |
| 4X | 1 | | B1 | 22.52 | 0.25 |
| | 2 | | B2 | 10.15 | 0.14 |
| | 3 | | B3 | 12.62 | 0.18 |
| | 4 | | B4 | 23.25 | 0.54 |
| | 5 | | B5 | 17.51 | 0.26 |
| | 6 | | B6 | 9.90 | 0.27 |
| | 7 | | B7 | 34.13 | 0.19 |
| | A | | A | 23.07 | 0.30 |
| | C | | C | 11.93 | 0.16 |
| | G | | G | 23.57 | 0.49 |
| | T | | T | 16.40 | 0.20 |
| 3X+Y | 2 | 3 | 2223 | 11.07 | 0.22 |
| | | | 2232 | 8.97 | 0.33 |
| | | | 2322 | 9.64 | 0.26 |
| | | | 3222 | 11.54 | 0.26 |
| | | 5 | 2225 | 11.16 | 0.49 |
| | | | 2252 | 11.36 | 0.13 |
| | | | 2522 | 11.19 | 0.22 |

| | | | | | |
|---|---|---|---|---|---|
| | | | 5222 | 11.48 | 0.35 |
| | | | 2333 | 9.25 | 0.16 |
| | | 2 | 3233 | 9.69 | 0.26 |
| | | | 3323 | 12.34 | 0.24 |
| | 3 | | 3332 | 12.64 | 0.39 |
| | | | 3335 | 13.36 | 0.16 |
| | | 5 | 3353 | 14.38 | 0.19 |
| | | | 3533 | 14.45 | 0.22 |
| | | | 5333 | 14.54 | 0.19 |
| | | | 2555 | 11.78 | 0.33 |
| | | 2 | 5255 | 11.48 | 0.45 |
| | | | 5525 | 15.31 | 0.22 |
| | 5 | | 5552 | 17.62 | 0.42 |
| | | | 3555 | 17.59 | 0.35 |
| | | 3 | 5355 | 16.02 | 0.19 |
| | | | 5535 | 15.82 | 0.21 |
| | | | 5553 | 17.14 | 0.28 |
| | | | 2323 | 9.65 | 0.15 |
| | | 3 | 2332 | 9.60 | 0.17 |
| **2X+2Y** | | | 3223 | 12.15 | 0.17 |
| | | | 3232 | 10.05 | 0.17 |
| | | 3 | 3322 | 13.66 | 0.18 |
| | 2 | | 2233 | 9.86 | 0.15 |
| | | | 2424 | 14.14 | 0.22 |
| | | | 2442 | 15.94 | 0.36 |
| | | 4 | 4224 | 14.57 | 0.17 |
| | | | 4242 | 15.22 | 0.18 |
| | | | 4422 | 15.80 | 0.35 |
| | | | 2244 | 12.58 | 0.15 |
| | 2 | | 2525 | 10.57 | 0.09 |
| | | | 2552 | 11.85 | 0.19 |
| | | 5 | 5225 | 12.15 | 0.10 |
| | | | 5252 | 11.55 | 0.09 |
| | | | 5522 | 14.48 | 0.19 |
| | | | 2255 | 11.19 | 0.16 |
| | 4 | | 4545 | 25.65 | 0.41 |

| | | | | | |
|---|---|---|---|---|---|
| | | 5 | | | |
| | | | 5454 | 20.99 | 0.38 |
| | | | 4554 | 22.27 | 0.28 |
| | | | 5445 | 20.74 | 0.45 |
| | | 5 | 5544 | 19.56 | 0.26 |
| | | | 4455 | 23.70 | 0.41 |
| | 1 | 2 | 1122 | 14.05 | 0.24 |
| | | 3 | 1133 | 18.93 | 0.21 |
| | | 4 | 1144 | 21.09 | 0.23 |
| | | 5 | 1155 | 20.50 | 0.25 |
| | 3 | 4 | 3344 | 20.18 | 0.87 |
| | | 5 | 3355 | 14.83 | 0.20 |

**Table S4**. The mean residual currents ($I_{res}$ (%)) and the full width half height values (FWHM) for each oligonucleotide were determined by performing Gaussian fits to the residual current histogram from experiments with different combination of natural and modified nucleotides at position x = 13 – 16 from the streptavidin anchor at 200 mV.

| Combination | X | Y | Sample | $I_{res}$ (%) | FWHM |
|---|---|---|---|---|---|
| **ACT+X** | 2 | | 2ACT | 12.08 | 0.33 |
| | | | ACT2 | 11.57 | 0.44 |
| | 5 | | 5ACT | 15.12 | 0.40 |
| | | | AC5T | 13.78 | 0.51 |
| | | | ACT5 | 12.26 | 0.30 |
| **4X** | 2 | | B2 | 12.08 | 0.11 |
| **3X+Y** | 2 | 3 | 2322 | 9.93 | 0.11 |
| | | 5 | 2225 | 11.60 | 0.61 |
| | | | 2252 | 12.59 | 0.09 |
| | | | 2522 | 12.21 | 0.16 |
| | | | 5222 | 13.25 | 0.10 |
| | 3 | 2 | 2333 | 10.00 | 0.16 |
| | | 5 | 3353 | 15.47 | 0.41 |
| | | | 3533 | 16.22 | 0.34 |
| | | | 5333 | 12.85 | 0.34 |
| | 5 | 2 | 2555 | 12.39 | 0.49 |
| | | | 5255 | 13.63 | 0.75 |
| **2X+2Y** | 2 | 3 | 2323 | 10.57 | 0.16 |
| | | | 2332 | 10.35 | 0.17 |
| | | | 3232 | 10.86 | 0.26 |
| | | 4 | 2442 | 17.19 | 0.25 |
| | | | 4422 | 17.98 | 0.55 |

#### The two-step event identification scheme for ONT readouts with NN processing

The main challenges faced when analyzing nanopore current signals are illustrated in **Figure S5.** The figure shows the extreme variations in the current levels, which can either stay close to the mean (as illustrated on the example CCCC) or deviate more than 15% from the mean (as illustrated on the example 2233). Therefore, to automatically extract the regions from the ONT current readouts that correspond to modified nucleotides without resorting to basecalling, we developed a two-step identification scheme depicted in **Figure S5.** The first step is to estimate the current level for the polyA region, which is subsequently used for calibration purposes. We used kernel density estimation of the signal level distribution (2), followed by identification of the levels that have the two largest probabilities in the estimated distribution. This approach is justified by the observation that in our oligo structure, the polyA regions constitute the longest signal component. As polyT current levels are expected to be lower than polyA levels, we subsequently filtered out readout regions that are trailed by nearly flat regions with a mean level value lower than that observed for the polyA tails, using a finite state machine (3).These regions are expected to bear the signal from the chemically modified nucleotides.

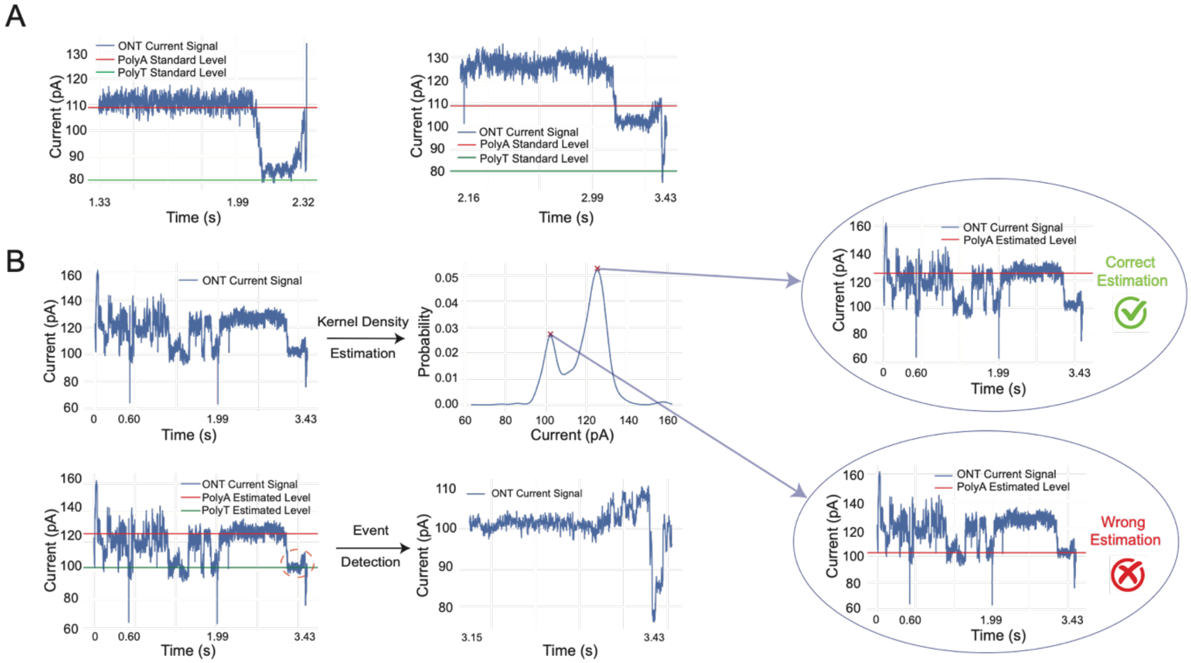


**Figure S5.** (**A**) (Left) Raw current readout of a control oligo bearing the content CCCC. (Right) A raw current readout bearing the content 2233. The red and green lines represent the expected standard levels for polyA and polyT regions, respectively. (**B**) Analysis of nanopore sequencing results for chemically modified nucleotides. (Top Left) Raw current readout for a control oligo containing the sequence 2233. (Top Right) Visualization of the kernel density estimation method: Two peaks correspond to two possible polyA region levels. (Bottom) The procedure for determining which level to use for calibration, based on the mean value of the "nearly-flat" region following the predicted polyA region. An example of the current level corresponding to the highest peak, which was used to correctly estimate the location of the polyA region. Building upon this step, the results show that one can also isolate the signal region which corresponds to the chemically modified nucleotides.

### Summary of the results of our model-based classification procedure

We trained ResNet models on 12 permutation classes in which the composition is fixed, but the orderings of the modified nucleotides are different. What we refer to as a "superclass" combines different choices and orderings of the modified nucleotides (the superclass contains 66 out of 77 tetramers, as for 11 tetramers an insufficient number of training samples was available). The number of valid sequenced reads (i.e., reads containing modified nucleotides) for each class is shown in **Table S5**. To perform unbiased training, we balanced out the sizes of the classes by setting a lower bound for subsampling of reads in different classes. We also set an upper bound on the number of training samples used for each class, in order to prohibit one/several classes to dominant the training set. For finer classification involving permutations of monomers within a class, we set the lower bound to 1000, and the upper bound to 5000. For the classification task on all 66 classes, we set the lower bound to 2000, and the upper bound to 3500. These choices are necessitated by two conflicting requirements: To balance out the class sizes and retain a training set as large as possible. The classification results are shown in **Figure S6**. From the confusion matrices we observe that almost all combinations can be easily distinguished from each other with very high accuracies (i.e., the diagonal values are significantly larger than the off-diagonal values). However, there are some tetramer instances that are hard to classify, such as 3223 (when compared to a tetramer in {2233, 3322, 2332, 3223, 2323, 3232}). The average classification accuracies for each model trained are listed in the caption of **Figure S6.**

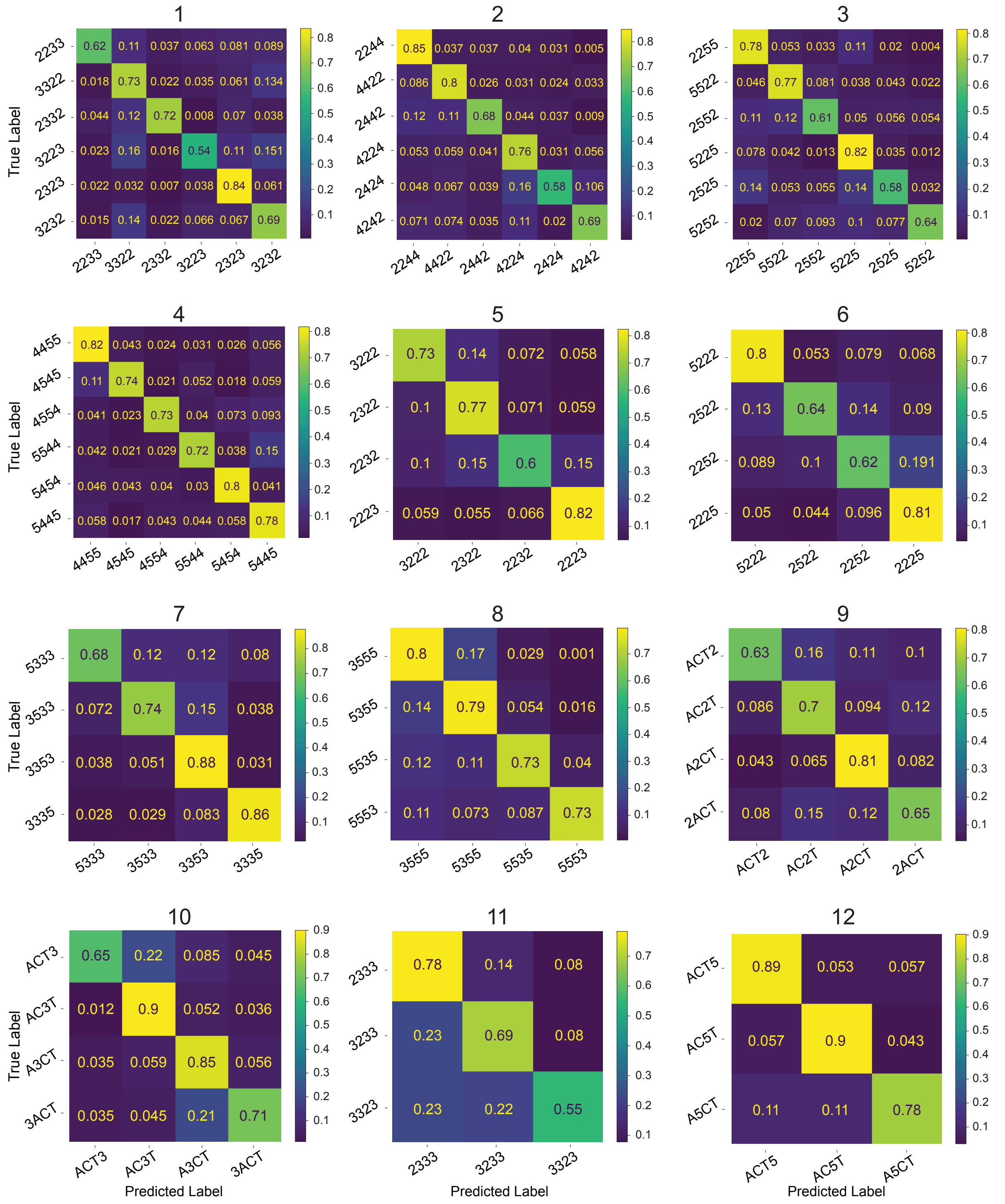


**Figure S6.** Classification performance of 12 different classes of tetramers. The names of the classes are listed in the subfigures, along with their average classification accuracies: (1) 69.39 ± 0.93%, (2) 72.25% ± 1.46%, (3) 68.87% ± 0.90%, (4) 77.84% ± 0.96%, (5) 72.18% ± 1.79%, (6)

71.97% ± 0.54%, (7) 81.27% ± 0.93%, (8) 79.17% ± 1.87%, (9) 69.66% ± 0.48%, (10) 80.04% ± 0.69%, (11) 70.81% ± 1.15%, (12) 88.00% ± 1.31%.

| Class Name | Number of valid reads | Class Name | Number of valid reads | Class Name | Number of valid reads | Class Name | Number of valid reads | Class Name | Number of valid reads |
|---|---|---|---|---|---|---|---|---|---|
| **3332** | 39 | **5255** | 74 | **2555** | 204 | **5ACT** | 315 | **7777** | 712 |
| **5525** | 750 | **TTTT** | 1390 | **ACT3** | 1717 | **3323** | 1808 | **3555** | 1885 |
| **5552** | 1944 | **A5CT** | 2133 | **5535** | 2315 | **3233** | 2344 | **5333** | 2430 |
| **5553** | 2460 | **4444** | 2553 | **GGGG** | 2607 | **6666** | 2632 | **2424** | 2706 |
| **1144** | 2723 | **4422** | 2740 | **1133** | 3134 | **3353** | 3167 | **4242** | 3310 |
| **4224** | 3377 | **3223** | 3732 | **ACT2** | 3837 | **3322** | 3865 | **2442** | 3967 |
| **2255** | 4039 | **4545** | 4072 | **4455** | 4500 | **3333** | 4506 | **5555** | 4630 |
| **5225** | 4657 | **4554** | 4827 | **2ACT** | 4827 | **1122** | 4844 | **5355** | 4925 |
| **A2CT** | 5197 | **CCCC** | 5198 | **5522** | 5236 | **3232** | 5324 | **3ACT** | 5403 |
| **5544** | 5485 | **AC2T** | 5505 | **2333** | 5612 | **5222** | 5905 | **2222** | 5958 |
| **5454** | 6090 | **5445** | 6163 | **3222** | 6395 | **2244** | 6484 | **2252** | 6509 |
| **3533** | 6526 | **AC5T** | 6532 | **3355** | 6556 | **2522** | 6799 | **2233** | 7047 |
| **2525** | 7403 | **A3CT** | 7448 | **2225** | 7563 | **1155** | 7591 | **2223** | 7700 |
| **3344** | 7716 | **AAAA** | 7927 | **3335** | 7952 | **2552** | 9525 | **2232** | 9955 |

| | | | | | | | | | |
|---|---|---|---|---|---|---|---|---|---|
| **ACT5** | 11768 | **1111** | 13502 | **2322** | 13915 | **2323** | 15927 | **5252** | 16104 |
| **2332** | 17890 | **AC3T** | 22040 | | | | | | |

**Table S5.** The number of valid reads for each tetramer class (77 classes in total), arranged in ascending order.
